# Reconstruction and identification of the native PLP synthase complex from Methanosarcina acetivorans lysate

**DOI:** 10.1101/2024.07.09.602819

**Authors:** Angela Agnew, Ethan Humm, Kang Zhou, Robert P. Gunsalus, Z. Hong Zhou

## Abstract

Many protein-protein interactions behave differently in biochemically purified forms as compared to their *in vivo* states. As such, determining native protein structures may elucidate structural states previously unknown for even well-characterized proteins. Here we apply the bottom-up structural proteomics method, *cryoID*, toward a model methanogenic archaeon. While they are keystone organisms in the global carbon cycle and active members of the human microbiome, there is a general lack of characterization of methanogen enzyme structure and function. Through the *cryoID* approach, we successfully reconstructed and identified the native *Methanosarcina acetivorans* pyridoxal 5’-phosphate (PLP) synthase (PdxS) complex directly from cryogenic electron microscopy (cryoEM) images of fractionated cellular lysate. We found that the native PdxS complex exists as a homo-dodecamer of PdxS subunits, and the previously proposed supracomplex containing both the synthase (PdxS) and glutaminase (PdxT) was not observed in cellular lysate. Our structure shows that the native PdxS monomer fashions a single 8α/8β TIM-barrel domain, surrounded by seven additional helices to mediate solvent and interface contacts. A density is present at the active site in the cryoEM map and is interpreted as ribose 5-phosphate. In addition to being the first reconstruction of the PdxS enzyme from a heterogeneous cellular sample, our results reveal a departure from previously published archaeal PdxS crystal structures, lacking the 37 amino acid insertion present in these prior cases. This study demonstrates the potential of applying the *cryoID* workflow to capture native structural states at atomic resolution for archaeal systems, for which traditional biochemical sample preparation is nontrivial.

## Importance

Archaea are one of the three domains of life, classified only recently as a phylogenetically distinct lineage (1). There is a paucity of known enzyme structures from organisms of this domain. This is further exacerbated by characteristically difficult growth conditions and a lack of readily available molecular biology toolkits to study proteins in archaeal cells. As a result, there is a gap in knowledge concerning the mechanisms governing archaeal protein behavior and their impacts on both the environment and human health; case in point, the synthesis of the widely utilized cofactor PLP (a vitamer of vitamin B6, which humans cannot produce). By leveraging the power of single particle cryoEM and map-to-primary sequence identification, we determine the native structure of PLP synthase from cellular lysate. Our workflow allows 1) rapid examination of new or less characterized systems with minimal sample, and 2) discovery of structural states inaccessible to recombinant expression.

## Introduction

While protein complexes from model eukaryotic and prokaryotic systems are highly represented in structural biology literature, there remain gaps in knowledge in cases of dynamic cellular processes and taxonomically neglected species. As members of one of the three domains of life alongside bacteria and eukaryotes, archaeal organisms have been chronically understudied due to several issues that have hindered accurate documentation of the domain. These include, but are not limited to, the incompatibility of bacterial identification strategies with archaeal cell structure, the sheer difficulty of culturing archaea, and the lack of a robust body of archaeal genome annotation in existing databases (2). Furthermore, many of these species can be recalcitrant to traditional molecular biology techniques, due to strict anaerobic growth conditions and a lack of established gene editing capabilities. While clonal and CRISPR-Cas9 techniques have been recently introduced for methanogenic archaeal clades, they are not currently commercially available nor widely distributed (3-5). However, often these unique cases provide crucial evolutionary context for foundational biology.

One such group of organisms are known as methanogens, a subset of the kingdom Euryarchaeota of the domain Archaea classified by their unique ability to anoxically metabolize organic compounds and produce methane. Methanogens are the primary source of biogenic methane on earth and are cornerstone to the anoxic carbon cycle. These archaea are ubiquitous, but mostly colonize habitats such as wetlands, sewage plants, and the stomachs of livestock (6). Methanogenesis is an ancient metabolism, with the first methanogen likely existing not long after bacteria and archaea diverged (7). We may already have an example of their tremendous influence on the biosphere, as the emergence of methanogens has been implicated in disrupting the global carbon cycle prior to a prehistoric era of mass extinction (8). Though still responsible for producing 1 gigaton of methane annually, much of the methane produced is subsequently metabolized by other methanotrophic microorganisms living in syntrophic association (6). The activity of methanotrophs is still insufficient to offset combined biogenic and non-biogenic sources of methane, such that the impact of microorganisms cannot be ignored in the effort to curtail greenhouse gas emissions.

Yet another feature of several methanogenic archaea is their relationship with the human microbiome. Archaeal species have now been identified from skin, the GI tract, and respiratory tract (9, 10). The consequence of cross-feeding with fermentative bacteria in this context is promoting overgrowth of pathogenic microbes, yet this field of research is still in its infancy. Still, there are some methanogenic archaea now tied to periodontitis, IBS, and colorectal cancer (11, 12). Since H_2_ is a methanogenic substrate, methanogens keep environmental concentrations of H_2_ low, energetically benefiting the fermentative metabolism of other bacteria. While these effects are not known to arise from virulence factors of the archaea themselves, these outcomes do result from their metabolic behavior and adherence to sites of infection. A mechanistic understanding of their key enzymes may give rise to therapeutic targets that can ameliorate these polymicrobial diseases.

Of current structural biology techniques, cryogenic electron microscopy (cryoEM) singularly enables us to achieve high resolution structures of large-scale, frozen-hydrated protein complexes in their native states (13). Previous publications have pioneered a pipeline that allows for autonomous reconstruction and identification of unknown proteins from images of an enriched cellular milieu, using an in-house software termed *cryoID* (14-16). By avoiding extensive genetic modifications, proteins may maintain the conformations, complexation, and chemical modifications crucial to their activity in living cells. To date, this method has not been applied to an archaeal system in pursuit of cryoEM reconstruction of native proteins.

Here, we have applied the *cryoID* approach to study the model methanogen species *Methanosarcina acetivorans*. By directly imaging and identifying native cytosolic proteins, we have successfully determined the PdxS subunit of its pyridoxal 5’-phosphate (PLP) synthase. Our reconstruction of the dodecamer reveals departures from the only two known crystal structures of Euryarchaeota Archaea and demonstrates the promise of the *cryoID* workflow for samples from less characterized systems, particularly those from Archaea, where molecular biology approaches have been limited.

## Results

### Overall Structure

In efforts to simultaneously ensure a sufficiently high population of dominant species in this sample while preserving as many native interactions as possible, per the *cryoID* approach, a glycerol density gradient alone was used to fractionate the sample by size. A 2mL fraction window containing particles 8-12 nm in diameter was pooled for structural examination due to the promising 2D class averages noted in negative stain electron microscopy (EM) screening (Figure 1B). Even after selecting a bespoke size range, resultant images of stained and frozen samples make evident the retained heterogeneity after separation (Figure 1A). Similar classes were noted between negative stain and cryoEM 2D averaging, and by employing a single particle analysis approach, we reconstructed a cryoEM map to 3.38 Å with D6 symmetry (Figure 1C). Even with just several views predominating, a high resolution map could be reconstructed. This made it rather feasible to “mix and match” various putative top and side views of different proteins in our 2D classes, and discover an *ab initio* reconstruction that produced a rational and continuous density. After repicking the dataset with a deep learning model trained on particles contributing to the best *ab initio* model, we successfully expanded our particle dataset selecting specifically for this bespoke species. Using the 3.4 Å resolution cryoEM reconstruction, we could identify the particle as the synthase (PdxS) subunit of the *Methanosarcina acetivorans* pyridoxal 5’ phosphate (PLP) synthase (previously shown to appear in supramolecular complex with its glutaminase subunit, PdxT) through the *cryoID* software, as well as model amino acids 11 to 301 of the monomeric subunit.

**Fig. 1.**
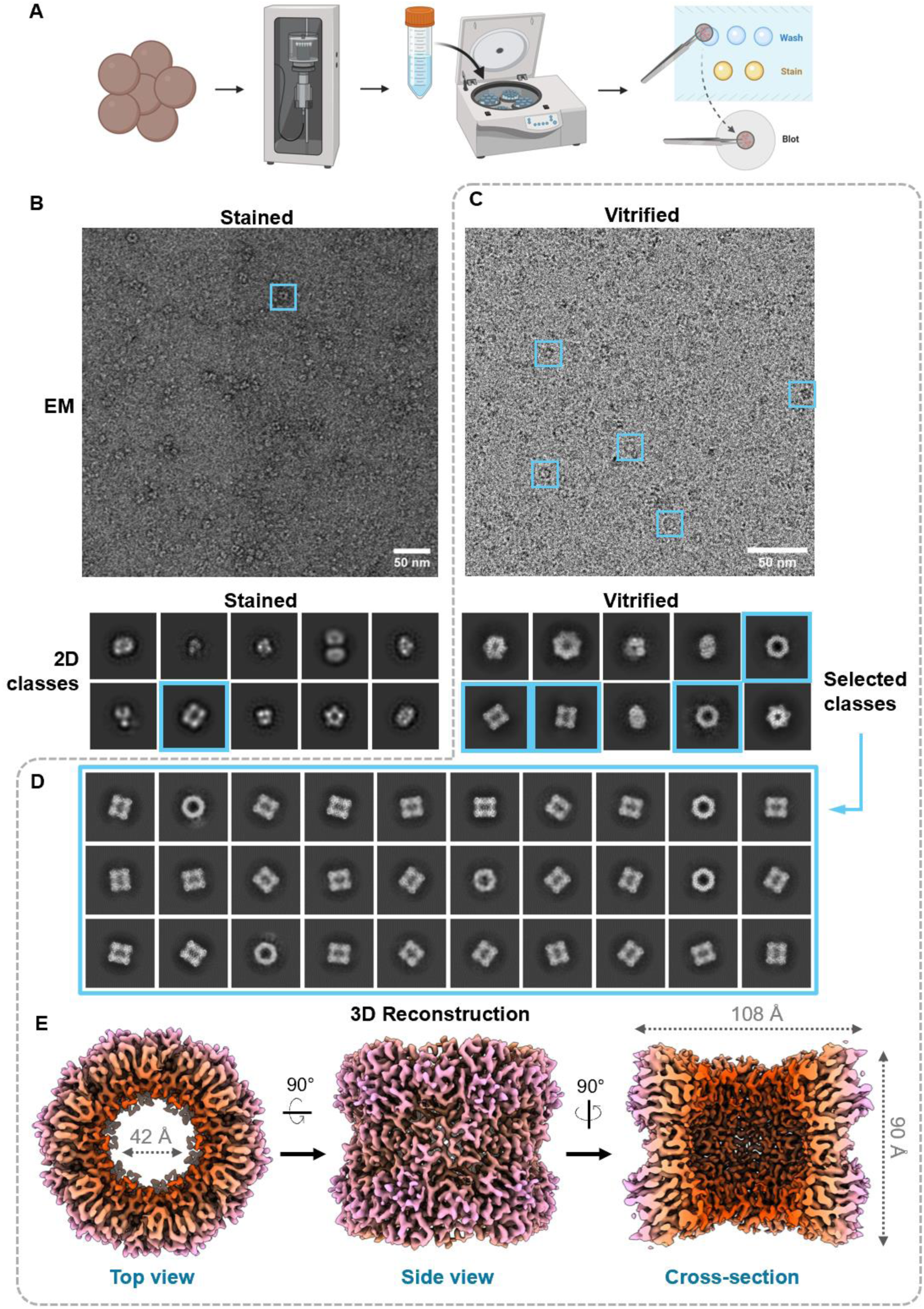
*CryoID* workflow for the reconstruction of PdxS. (*A*) *M. acetivorans* cells are lysed by sonication, fractionated by size, and applied to TEM grids. (*B*) Representative negative stain TEM micrograph of selected density gradient fraction and corresponding 2D class averages. An example PdxS particle and its corresponding 2D class are both boxed in blue. (*C-E*) Cryo data reconstruction workflow (boxed in dotted gray). An example cryoEM micrograph (*C*) illustrates the extent of sample heterogeneity, with particles of interest corresponding to PdxS boxed in blue. Like in panel *B*, corresponding 2D class averages are boxed in blue. After selecting initial classes that represent top and bottom views, the dataset was repicked with a Topaz model trained on the selected classes, giving rise to the second set of 2D classes exclusively showing different views of our chosen particle (*D*). Using D6 symmetric nonhomogenous reconstruction in cryoSPARC, we obtain the map seen in panel *E*. Density map is colored by cylindrical radius.

Like other homologous PLP synthase structures, the *M. acetivorans* PdxS dodecameric complex consists of two layers of homohexameric rings that stack cylindrically, with an outer diameter of 108 Å and an inner diameter of 42 Å. Each monomer adapts a triose phosphate isomerase (TIM) barrel fold that houses the active site for deoxyxylulose 5-phosphate (DXP)- independent PLP synthesis from D-ribose 5-phosphate (R5P), D-glyceraldehyde 3-phosphate (G3P), and ammonia. Ribulose 5-phosphate (Ru5P) and dihydroxyacetone phosphate (DHAP) are also acceptable substrates, as PdxS demonstrates triose and pentose isomerase activity in addition to PLP synthesis (as suggested by the domain architecture) (17). There is strong density in this binding pocket adjacent to residues annotated as binding sites for R5P, suggestive of captured particles undergoing native catalysis. The first ten residues lack clear density for modeling, as was similarly observed with the *Arabidopsis thaliana* Pdx1.2/1.3 pseudoenzyme structures (18), indicating that this region is relatively flexible when unbound to its glutaminase, PdxT. The rest of the monomeric secondary structure elements are named by sequence, as depicted in Figure 1B. There are in total 15 alpha helices and 8 beta sheets, with helices α1-8 (excluding prime and double-prime alpha helices) participating in the TIM-barrel fold with beta sheets β1-β8.

### Interactions between PdxS subunits

The atomic model of the PdxS complex shows high levels of surface and charge complementarity in its inter-subunit binding interfaces (Figure 3A, 3D). The monomer itself resembles an asymmetric, inverted triangle, demonstrating on either side a convex (A) or concave (B) facet posed for lateral binding. Monomer tips also interdigitate at the hexamer-hexamer interface, creating altogether three distinct types of lateral contacts that give rise to the dodecamer, here described as AB, AA, and BB (Figure 3B). A network of 6 inter-subunit hydrogen bonds anneal the facets of interface AB together, while 6 such interactions comprise AA, and another 6 comprise BB (while each have 3 unique bonds respectively, these are doubled due to the inherent two-fold symmetry of the “self” interfaces). The residues contributing to the AB contacts are as follows. One hydrogen bond forms between glutamate 98 of helix 3 and arginine 233 of helix 7. Also posed on α7 is aspartate 227, which forms two hydrogen bonds; one with histidine 93 from loop β3 to α3, and arginine 90 on β3. Arginine 67 from loop α2’-α2 forms a backbone contact with methionine 282 of loop α8”-α9. Loop α8”-α9 also supports a glycine backbone H-bond to glutamate 72 on α2. Lastly, aspartate 118 from loop β4 to α4 forms a hydrogen bond with lysine 172 of α6. The “self” interfaces (AA and BB) are dominated by electrostatic interactions between alpha helices 6 and 6’. Comprising the AA interface, the three unique sidechain bond pairs include hydrogen bonds between lysine 186 (α6’) to aspartate 124 (α6-6’), arginine 194 (α6’) to glutamate 197 (α6’), and arginine 194 to the C-terminal carboxylate of tryptophan 301. For the contacts involved in BB, the three unique bonds include a hydrogen bond between lysine 172 (α6) and glutamate 187 (α6’), and another two between arginine 179 (α6) and glutamate 188 (α6’).

**Fig. 2.**
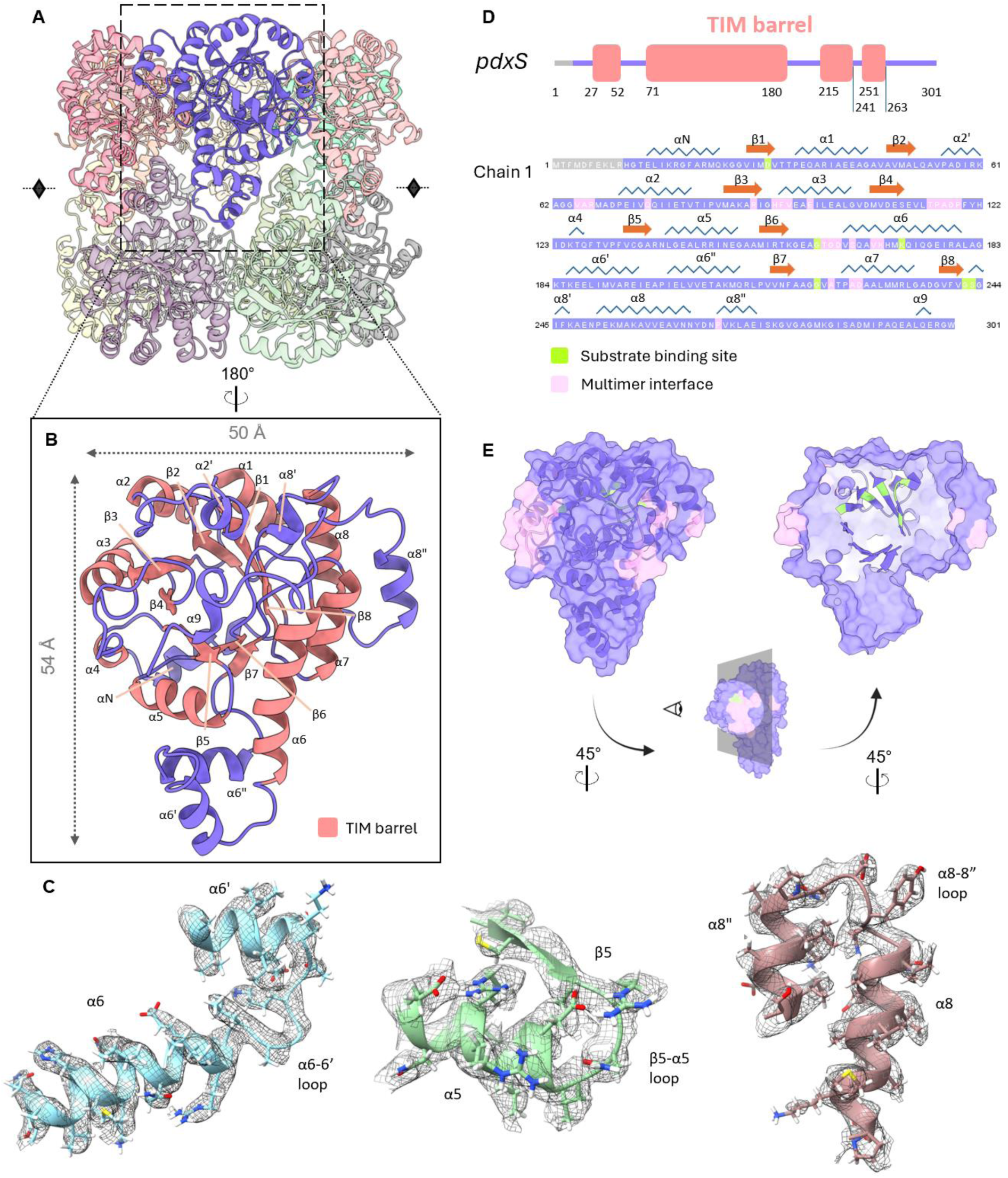
Monomer structure of PdxS. (*A*) Multimer structure of PdxS with one subunit highlighted. The monomer subunit is depicted as a ribbon representation with secondary structure elements labeled (*B*) and with selected sections docked into the cryoEM density (represented as a mesh) to demonstrate sidechain fit (*C*). (*D*) Domain structure of the PdxS monomer and its complete primary sequence with secondary structure elements, binding site residues, and multimer interface interactions annotated according to UniProt entry (42). (*E*) Space-filling representation of the monomer model with multimer interfaces colored in pink and R5P binding site residues colored in green, along with a second cut-away view to reveal the beta barrel ribbon depiction within.

**Fig. 3.**
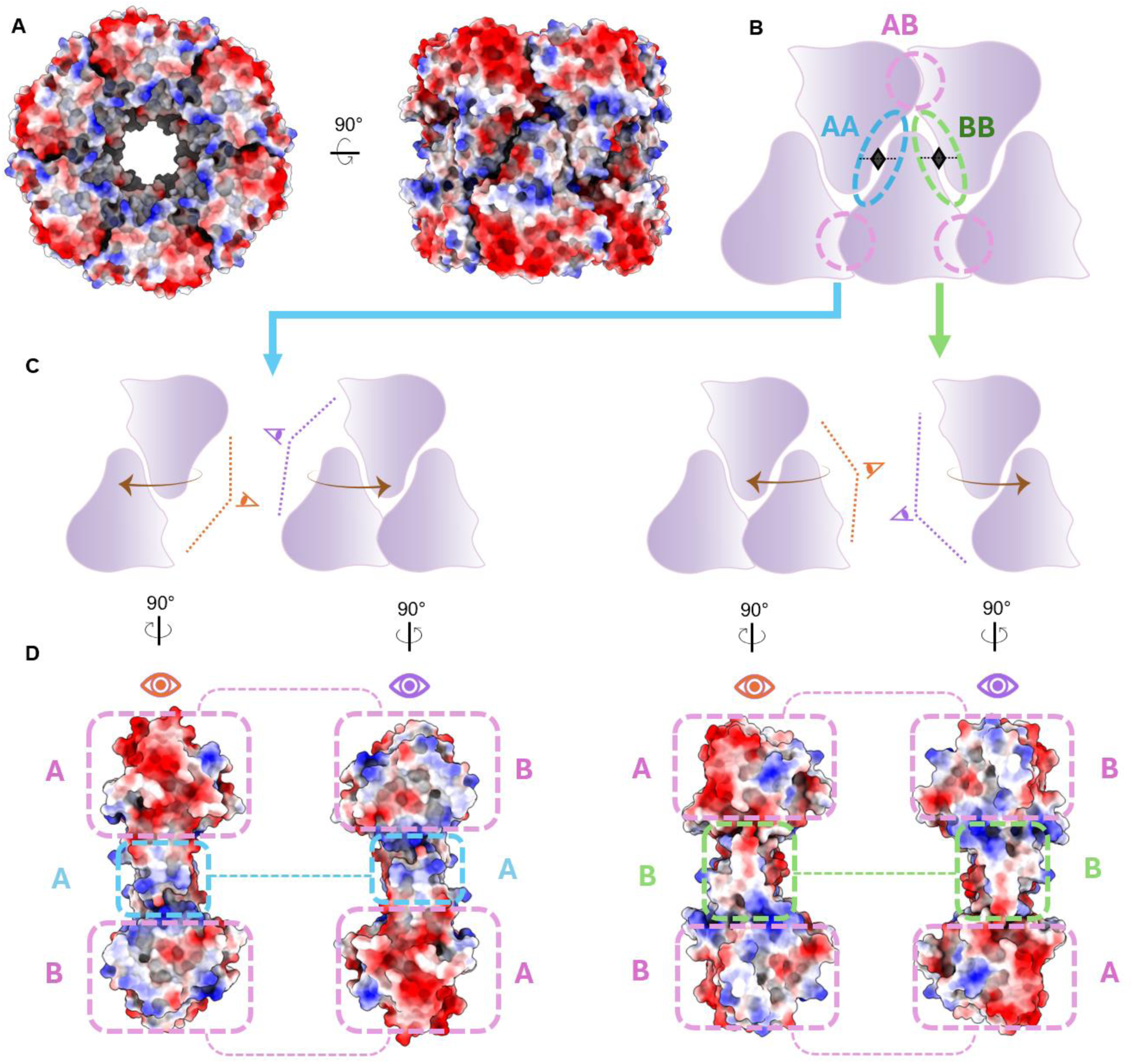
Charge and shape complementarity in oligomeric interfaces. (*A*) Top and side view of the space-filling representation of the PdxS dodecamer with surface electrostatics depicted in red (acidic) and blue (basic). (*B*) Cartoon schematic of intersubunit interactions defining each of three types of contacts, based on splitting the asymmetric monomer unit into two unique lateral faces, A and B. Each contact region is assigned a corresponding annotation color. (*C*) Illustration of views for each interface to be depicted in panel *D*. (*D*) Electrostatic surface coloring of complementary interfaces with each unique contact region annotated in its corresponding color as defined in (*B*). The AA and BB regions are self-contacts. Overall, acidic and basic patches along each surface are complementary.

The overall shape of the monomer is topologically complementary to itself, given that monomers within a hexamer self-associate via the concave-convex fit of the A and B interfaces. The alpha helices external to the (β/α)_8_ TIM barrel motif (αN, α2’, α6’, α6”, α8’, α8”, and α9) are also responsible for some morphological stabilization in addition to protecting the active site from solvent exposure (αN and α2’). These stabilizing contacts include a protrusion of the T115-Y121 loop into a hydrophobic patch along the crevice between the adjacent monomer’s α6 and α7 helices along the AB interface. Another such region is located at the same interface, anterior to the aforementioned loop; α7 and α8" share valine and (iso)leucine-mediated contacts with the adjacent monomeric unit’s α3 helix.

In our maps, no instances of PdxT binding were observed, even after extensive 3D classification and examination under a low-density threshold. Furthermore, upon screening the higher molecular weight fractions off of the glycerol density gradient, no abundance of PdxS-PdxT supracomplexes were present as determined by 2D classification and examination of individual micrographs. In homologous proteins for which there are crystal structures, the αN helix of PdxS is instrumental to PdxT binding (19). The *M. acetivorans* αN helix displays several solvent-exposed hydrophobic residues (I16, G19, F20, M23), with regions of dense negative and positive charge above and below, respectively, which is highly suggestive of a putative binding interface for the glutaminase. That being said, given that this sample represents a composite of the most dominant states in our cellular lysate, our work may suggest a novel state wherein PdxT binds extremely transiently and upon delivery of substrates, need not remain docked to the PdxS homo-dodecamer.

### Active site chemistry and evolutionary conservation

On the interior of the TIM barrel, we observe an elongated density adjacent to residues annotated as substrate binding sites (Figure 4A). The PLP synthase active site chemistry as currently understood is rather complex, involving ring openings, closings, and isomerizations, but generally proceeds with a linearized R5P being stabilized via formation of a Schiff base of its C1 with an active site lysine residue (19). A secondary lysine residue likewise bonds with the C5 of R5P, and is eventually responsible for swinging the intermediate to a second phosphate binding site exterior to the TIM barrel (20). Based on comparison with previous crystal structures where PdxS was co-crystallized with R5P and various intermediate states, the density is most likely to belong to either bound R5P or a R5P Schiff-base intermediate. Docking a R5P moiety with the molecule oriented such that the phosphate group is positioned in the bulbous density nestled between β6 and loop 156-164 then positions the oxygen 1 of R5P next to lysine 88, which is where the structurally equivalent lysine in other structures (e.g. *Plasmodium berghei*) forms a Schiff base intermediate with R5P (Figure 4B-E). This also positions R5P to engage in a hydrogen bond with aspartate 31. One caveat to this assignment is that the oblong density adjacent to the putative phosphate density does, however, run in a perpendicular direction to the conserved R5P position. Repositioning the R5P to align with this perpendicular density could suggest engagement of the second active site lysine (K156) in formation of a second Schiff base with C5, but the density in our map for K156 is very strongly oriented in its alternate conformation facing away from the TIM barrel active site (Figure 4C). It is clear, therefore, that multiple substrate states have likely been captured in this averaged density map.

**Fig. 4.**
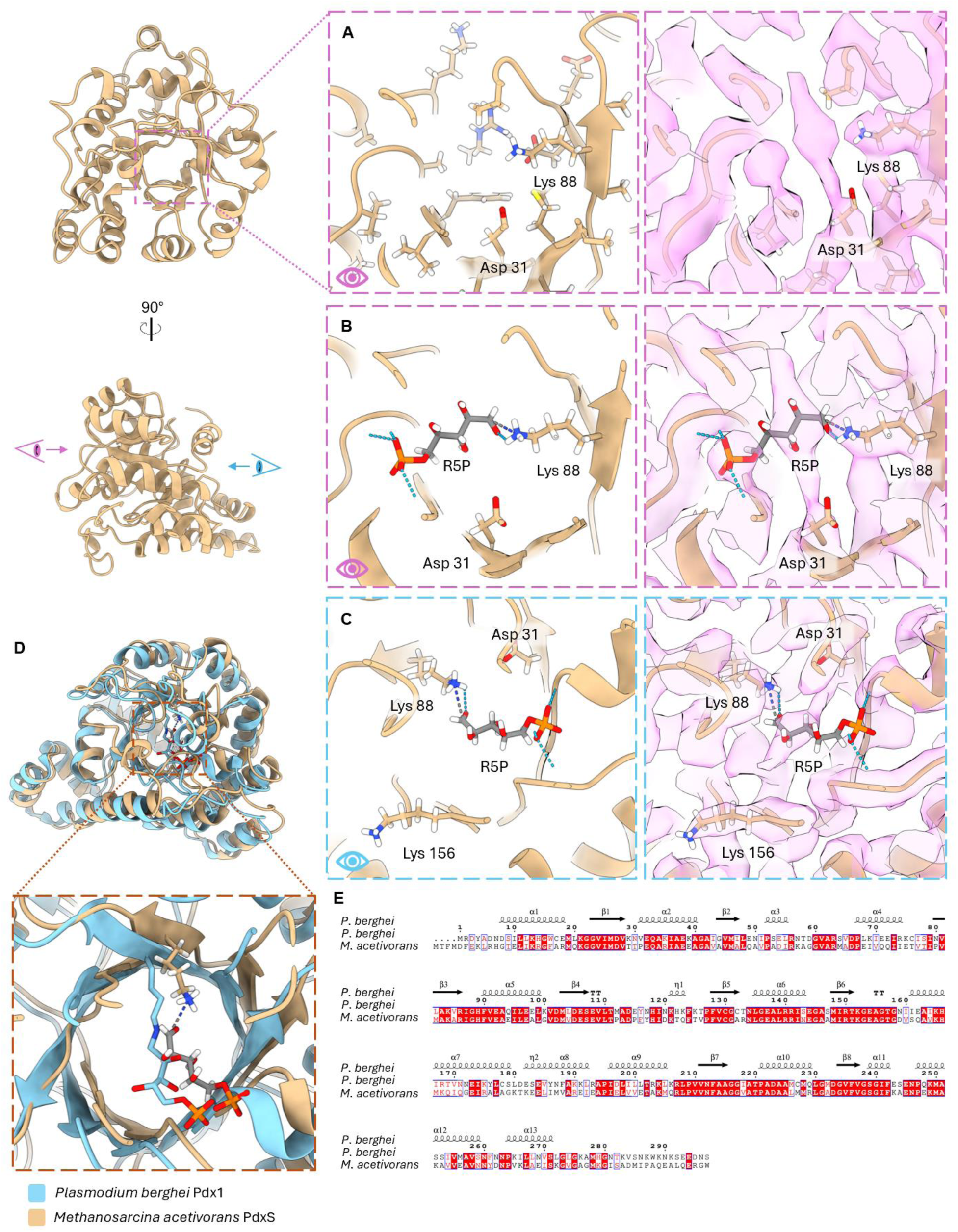
Active site densities and potential ligand modeling. (*A*) Ribbon model and corresponding CryoEM density of PdxS in its active site. Nearby active site residues are represented as stick models. The elongated density and its neighboring spherical density colored in pink are not occupied by any nearby side chains. (*B-C*) Possible R5P configuration in active site density, with the phosphate group docked into the spherical density. A potential Schiff base is represented with K88 as a dashed dark blue and grey line. Hydrogen bonds between the phosphate group and nearby backbone are depicted with a dashed light blue line. (*C*) Alternative view of the docked R5P, revealing the strong density for the second active sight lysine (K156) in its extra-active site conformation. (*D-E*) Comparison of *Plasmodium berghei* and *Methanosarcina acetivorans* Pdx structure (*D*) and primary sequence (*E*). The crystal structure for *P. berghei* bound with R5P is aligned with our structure, including our hypothesized R5P placement in the density.

Upon performing a phylogenetic analysis of the PLP synthase sequences from other biological systems for which crystal structures have been deposited, it is apparent that the *M. acetivorans* PdxS shares more sequence and structural homology with the other deposited bacterial and eukaryotic PLP synthases than with the two archaeal systems so far determined (*Pyrococcus horikoshii* and *Methanococcus jannaschii*) (21, 22). This is due to a 37-aa insertion that appears in various archaeal classes within the kingdom Euryarchaeota (a kingdom shared with *M. acetivorans*), though notably this mainly appears within extremophile genera. The insertion gives rise to an extra alpha helix and beta sheet between equivalent helices to *M. acetivorans* α6’ and α6”.

## Discussion

In this study, we have determined the first native structure of the PdxS component of PLP synthase from an archaeon by cryoEM and the *cryoID* approach. We demonstrate that the high degree of structural and sequence conservation witnessed in other crystal structures across bacteria and eukaryotes, is also shared in this archaeal lineage. Intriguingly, our model shows greater divergence from the sole deposited archaeal PLP synthase structures at current than all other taxa for which the synthase structure is known. Given that these two species are extremophiles, the helical insertions may provide greater stability of the oligomeric interfaces under high temperature conditions. The comparisons also shed light upon early evolutionary history, as this could further evidence the crosstalk between diverging archaeal and bacterial lineages via horizontal gene transfer, resulting in highly conserved properties such as *de novo* PLP synthesis coexisting with highly divergent properties such as s-layer and membrane composition (23-26).

Pyridoxal 5’-phosphate is a vitamer of vitamin B6, which is an essential cofactor for human neurological and immune health yet is one that humans and animals cannot natively synthesize. PLP is the biologically active state of vitamin B6, participating in over 160 crucial enzymatic processes which include the metabolism of glycogen, amino acids, and lipids (Figure 5) (27, 28). Particularly due to its roles in production of neurotransmitters and modulation of interleukin-2, PLP deficiencies have been correlated with neurological disorders such as epileptic encephalopathy, schizophrenia, and Parkinson’s disease, as well as immune system dysregulation and heightened inflammation (29, 30). Since humans can only obtain vitamin B6 from their diet or from endogenous microbial species, it is clear that microbiome composition and activity have an effect on vitamin B6 utilization (31). As components of the gut and oral microbiome, vitamin B6 production may represent another route through which archaea have specific relevance to human health outcomes, as our work both confirms the expression and particular abundance of PLP synthesis in our model species *M. acetivorans*. Furthermore, the fact that humans lack the genes for *de novo* vitamin B6 synthesis suggests that PLP synthases are a potentially druggable target for microbial disease. Some methanogenic archaea can grow mutualistically with environmental bacterial colonies, marking them for examination in the development of therapeutics against pathogenic bacterial species.

**Fig. 5.**
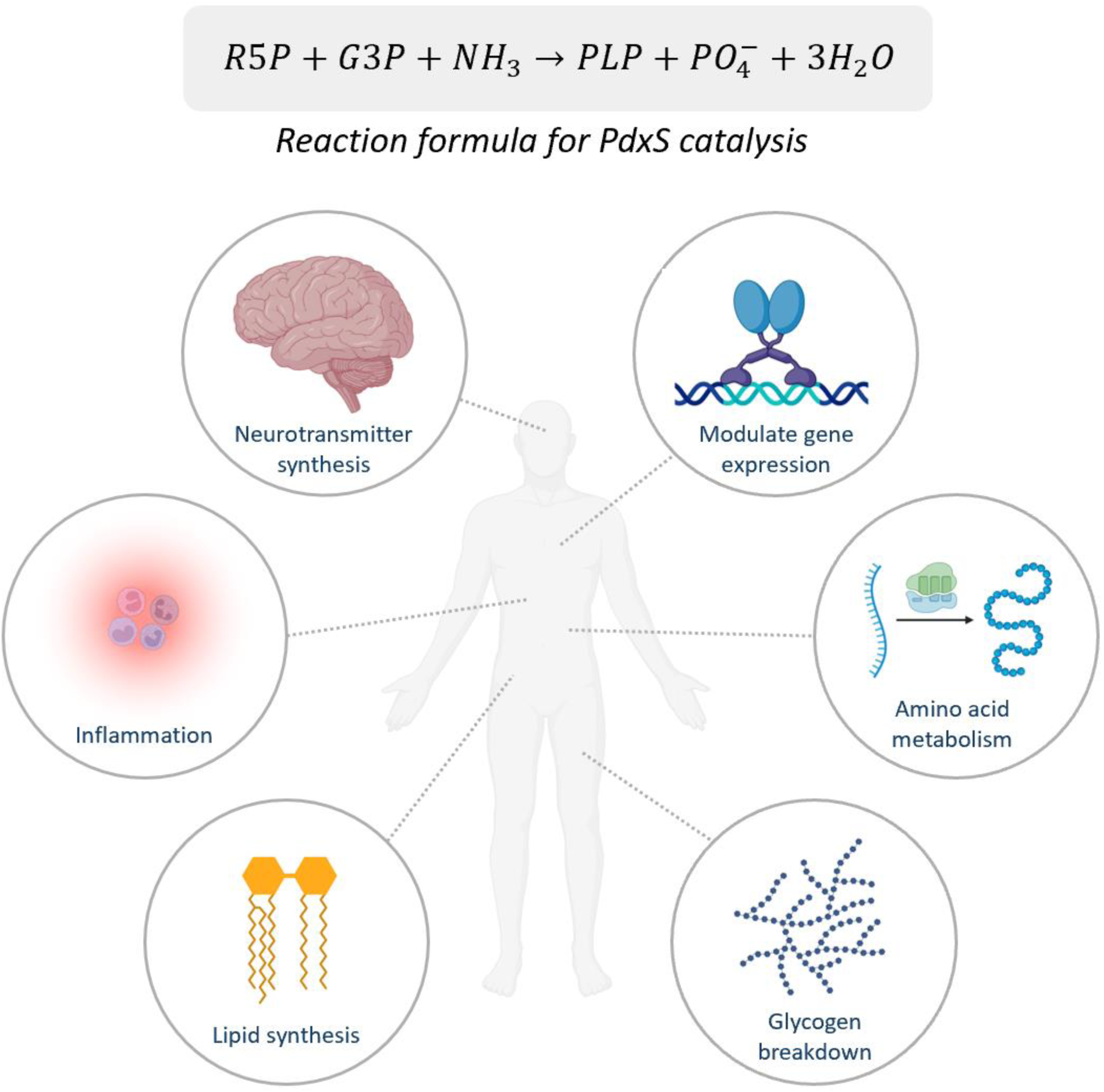
PLP is an essential cofactor for many basic cellular processes implicated in human health, including biosynthesis and metabolism of amino acids, carbohydrates, lipids, and nucleic acids; neurotransmitter production; modulating steroidal receptor gene expression; and regulating immune function and inflammatory responses (17, 43, 44).

Our present work also demonstrates the promise of leveraging the *cryoID* workflow for native proteins harvested directly from cellular lysate. With only crude size-based fractionation, current single particle analysis capabilities were successful in selecting, reconstructing, and identifying a native protein. Furthermore, we observe in our density some evidence of active site occupancy. Because of the multiple orientations of R5P or R5P intermediates this density accommodates, it is likely that we captured an amalgamation of many catalytic states. With further imaging and a greater particle set, these states can be classified and separately reconstructed. As such, this particular promise is fruitful for studying native interactions involved in catalytic cycles of enzymes, or for those systems which lack developed or established molecular biology techniques for extensive gene editing and endogenous protein purification. Structural characterization of biomolecules is crucial to 1) the development of targeted therapies in disease-causing systems, and 2) the pursuit of evolutionary descriptions of function. This shows it is essential to leverage the gross information retrieval capabilities of bottom-up cryoEM toward understudied organisms, such as those in archaeal taxa.

## Materials and Methods

### Sample Preparation

*M. acetivorans* C2A (DSM 2834) was cultivated in 100 mL serum bottles with a N2:CO2 (80:20) headspace and 50 mL medium as previously described (32). Methanol was used as the sole carbon and energy supply (50 mM). Cells were harvested at OD ∼ 1 by centrifugation at 10,000 x g for ten minutes at 5°C in an IEC tabletop centrifuge. Cells were resuspended in chilled lysis buffer (137 mM NaCl, 2.7 mM KCl, 10 mM Na2HPO4, 1.8 mM KH2PO4, 5 mM MgCl, 2 mM dithiothreitol (DTT), and protease inhibitor) and lysed via 1.5-150 Watt adjustable sonication at 20-25 kHz (40% power using ten 2s pulses at 4°C). Cell lysate was incubated with benzonase for 10 min before centrifugation (12,000 g for 6 min at 4°C) to clarify nucleic acid and cellular debris. The supernatant was decanted, applied to a 10-30% glycerol density gradient, and centrifuged at 110,000 g for 20 hours at 4°C. 500 µL gradient fractions were evaluated by SDS-PAGE and negative stain electron microscopy (EM), and selected fractions containing particles 8-12 nm in diameter were pooled together for subsequent grid preparation.

### Electron microscopy (EM) of stained and vitrified samples

For negative stain EM screening, 3 µL aliquots were applied to glow-discharged formvar/carbon-coated grids (300 mesh, Ted Pella) and incubated for 1 min, before negative staining with 2% uranyl acetate. Samples were screened on an FEI Tecnai F20 electron microscope operated at 200 keV, and images were recorded on a TIETZ F415MP 16-megapixel CCD camera at a nominal magnification of 60,000×.

Prior to freezing, n-dodecyl-beta-maltoside (DDM) was added to the sample to a final concentration of 0.0043% (one half the CMC) to alleviate protein denaturation arising from aggregation at the air-water interface. Subsequently, 3 µL aliquots were applied to glow-discharged holey carbon grids (Quantifoil R1.2/1.3 300 mesh, Ted Pella) and incubated for 10 s before automated blotting and flash-freezing in liquid ethane with a Vitrobot Mark IV vitrification system (Thermo Fisher Scientific). Various freezing conditions—including chamber temperature, humidity, blotting time, blotting force, and drain time after blotting—were screened on the aforementioned instrument used during negative stain evaluation. Optimal conditions were obtained with a 90 s glow discharge, a chamber temperature of 4°C and 100% humidity, 5 s blotting time, blot force of 0, and 0 s drain time. Optimized cryoEM grids were stored in liquid nitrogen until cryoEM data collection.

Imaging for downstream processing was performed on a Titan Krios 300 kV electron microscope (Thermo Fisher Scientific) equipped with a Gatan Imaging Filter (GIF) Quantum LS and a Gatan K3 direct electron detector. 14,890 movies were recorded with SerialEM in super-resolution mode at a nominal magnification of 81,000×, yielding a calibrated pixel size of 0.55 Å/pixel at the specimen level (33). The GIF slit width was set to 20 eV. Each movie contained 44 frames with an exposure time of 2.2 s per frame, giving a total estimated electron dose of 50 e-/ Å2.

### Structure determination

Movie frames were aligned with patch motion correction in the cryoSPARC v4 suite, after which the calibrated pixel size was 1.1 Å/pixel (34). Defocus values were determined with the cryoSPARC patch CTF estimation job. Autonomous, reference-free particle picking was first performed with the Blob Picker tool, specifically selecting for particles in the size range of 80-120 Å diameter. Initially 14,627,499 particles were picked from 14,890 micrographs, and boxed out by 300 × 300 pixels. After several iterations of 2D classification to remove junk classes, three distinct classes demonstrating alignment to 6 Å resolution emerged. These classes were used as templates to repick the dataset, which after iterative 2D classification yielded 17 well-aligned classes of 614,949 particles. This particle subset was subsequently used to train the deep learning-based particle picker Topaz, after which the entire dataset was again repicked with the trained model, yielding 3,724,083 particles (35). Another round of iterative 2D classification gave rise to many well-aligned, distinct 2D classes. Of these, several classes appeared to represent a 100 Å D6-symmetric structure. These were used for training of a separate Topaz model, and the dataset was picked a third time with a model trained on just these views. From the resultant 2D classes, a total of 99,971 particles were selected and subjected to ab initio reconstruction in cryoSPARC.

This initial model was used as a reference for nonuniform refinement, and with D6 symmetry enforced, the map reconstructed to 3.59 Å. After 3D classification and iterative CTF refinement and 3D refinement, the final resolution was 3.38 Å based on the gold standard FSC 0.143 criterion. B-factor, local resolution, and FSC curves were all calculated in cryoSPARC.

### Structure Identification with CryoID

After feeding *cryoID* our sharpened map, the program generated two query sequences built into two helical domains. The model was extended on both termini as permitted by the density, resulting in the following query sequences: 1) GGKYLKLPLGLGKGLGGKGLL, and 2) LPGKLGLGG. The top ranked output was UniProt Q8TQH6, or PdxS, which was further validated by docking the Alphafold3-predicted structure into our density.

### Atomic Modeling, Model Refinement and Graphics Visualization

Modeling was performed by docking the Alphafold3 predicted structure of the PdxS monomer into the cryoEM density, and confirming that the output identification was rational. The model was then iteratively real-space refined in the ISOLDE package of UCSF ChimeraX and validated in PHENIX, until outlier scores became negligible (36-38). A last round of real space refinement was performed in PHENIX, and evaluated using the PDB validation server (39). Visualization of all maps and models was performed with UCSF ChimeraX. All sequence alignments were performed with ClustalW and visualized with ESPript 3 (40, 41).

## ACKNOWLEDGEMENT

This project is supported partly by grants from the US NIH (R01GM071940 to Z.H.Z.) and the National Science Foundation (1911781 to R.P.G.) and U.S. Department of Energy (DOE) Office of Science (BER) (contract DE-FC-02-02ER63421 to R.P.G.). We acknowledge the use of instruments at the Electron Imaging Center for Nanomachines supported by UCLA and grants from the NIH (1S10OD018111) and the National Science Foundation (DBI-1338135 and DMR-1548924). Research reported in this publication was supported by the National Institute of General Medical Sciences of the National Institutes of Health under Award Number T32GM145388. The content is solely the responsibility of the authors and does not necessarily represent the official views of the National Institutes of Health. We thank David Strugatsky for his assistance in cryoEM data collection.

## Author contributions

Z.H.Z. and R.P.G. conceived the project; R.P.G., E.H., and A.A. prepared samples; A.A. and K.Z. recorded cryoEM images, A.A. processed the data and built the atomic model; A.A., Z.H.Z., and R.P.G. interpreted results and wrote the paper; all authors edited and approved the paper.

## References

1. Woese CR, Kandler O, Wheelis ML. Towards a natural system of organisms: proposal for the domains Archaea, Bacteria, and Eucarya. Proc Natl Acad Sci U S A. 1990;87(12):4576–9. Epub 1990/06/01. doi: 10.1073/pnas.87.12.4576. PubMed PMID: 2112744; PMCID: PMC54159.

2. Koskinen K, Pausan MR, Perras AK, Beck M, Bang C, Mora M, Schilhabel A, Schmitz R, Moissl-Eichinger C. First Insights into the Diverse Human Archaeome: Specific Detection of Archaea in the Gastrointestinal Tract, Lung, and Nose and on Skin. mBio. 2017;8(6). Epub 2017/11/16. doi: 10.1128/mBio.00824-17. PubMed PMID: 29138298; PMCID: PMC5686531.

3. Metcalf WW, Zhang JK, Apolinario E, Sowers KR, Wolfe RS. A genetic system for Archaea of the genus Methanosarcina: liposome-mediated transformation and construction of shuttle vectors. Proc Natl Acad Sci U S A. 1997;94(6):2626–31. Epub 1997/03/18. doi: 10.1073/pnas.94.6.2626. PubMed PMID: 9122246; PMCID: PMC20139.

4. Nayak DD, Metcalf WW. Cas9-mediated genome editing in the methanogenic archaeon Methanosarcina acetivorans. Proc Natl Acad Sci U S A. 2017;114(11):2976–81. Epub 2017/03/08. doi: 10.1073/pnas.1618596114. PubMed PMID: 28265068; PMCID: PMC5358397.

5. Kohler PR, Metcalf WW. Genetic manipulation of Methanosarcina spp. Front Microbiol. 2012;3:259. Epub 2012/07/28. doi: 10.3389/fmicb.2012.00259. PubMed PMID: 22837755; PMCID: PMC3403347.

6. Thauer RK, Kaster AK, Seedorf H, Buckel W, Hedderich R. Methanogenic archaea: ecologically relevant differences in energy conservation. Nat Rev Microbiol. 2008;6(8):579–91. Epub 2008/07/01. doi: 10.1038/nrmicro1931. PubMed PMID: 18587410.

7. Wang Y, Wegener G, Williams TA, Xie R, Hou J, Tian C, Zhang Y, Wang F, Xiao X. A methylotrophic origin of methanogenesis and early divergence of anaerobic multicarbon alkane metabolism. Sci Adv. 2021;7(27). Epub 2021/07/04. doi: 10.1126/sciadv.abj1453. PubMed PMID: 34215592.

8. Rothman DH, Fournier GP, French KL, Alm EJ, Boyle EA, Cao C, Summons RE. Methanogenic burst in the end-Permian carbon cycle. Proc Natl Acad Sci U S A. 2014;111(15):5462–7. Epub 2014/04/08. doi: 10.1073/pnas.1318106111. PubMed PMID: 24706773; PMCID: PMC3992638.

9. Borrel G, Brugere JF, Gribaldo S, Schmitz RA, Moissl-Eichinger C. The host-associated archaeome. Nat Rev Microbiol. 2020;18(11):622-36. Epub 2020/07/22. doi: 10.1038/s41579-020-0407-y. PubMed PMID: 32690877.

10. Candeliere F, Sola L, Raimondi S, Rossi M, Amaretti A. Good and bad dispositions between archaea and bacteria in the human gut: New insights from metagenomic survey and co-occurrence analysis. Synth Syst Biotechnol. 2024;9(1):88–98. Epub 2024/01/31. doi: 10.1016/j.synbio.2023.12.007. PubMed PMID: 38292760; PMCID: PMC10824687.

11. Cai M, Tang X. Human Archaea and Associated Metabolites in Health and Disease. Biochemistry. 2022;61(24):2835–40. Epub 2022/07/01. doi: 10.1021/acs.biochem.2c00232. PubMed PMID: 35770746.

12. Coker OO, Wu WKK, Wong SH, Sung JJY, Yu J. Altered Gut Archaea Composition and Interaction With Bacteria Are Associated With Colorectal Cancer. Gastroenterology. 2020;159(4):1459–70 e5. Epub 2020/06/23. doi: 10.1053/j.gastro.2020.06.042. PubMed PMID: 32569776.

13. Saibil HR. Cryo-EM in molecular and cellular biology. Mol Cell. 2022;82(2):274–84. Epub 2022/01/23. doi: 10.1016/j.molcel.2021.12.016. PubMed PMID: 35063096.

14. Ho CM, Li X, Lai M, Terwilliger TC, Beck JR, Wohlschlegel J, Goldberg DE, Fitzpatrick AWP, Zhou ZH. Bottom-up structural proteomics: cryoEM of protein complexes enriched from the cellular milieu. Nat Methods. 2020;17(1):79–85. Epub 2019/11/27. doi: 10.1038/s41592-019-0637-y. PubMed PMID: 31768063; PMCID: PMC7494424.

15. Xia X, Liu S, Zhou ZH. Structure, dynamics and assembly of the ankyrin complex on human red blood cell membrane. Nat Struct Mol Biol. 2022;29(7):698–705. Epub 2022/06/03. doi: 10.1038/s41594-022-00779-7. PubMed PMID: 35655099; PMCID: PMC9489475.

16. Liu S, Xia X, Calvo E, Zhou ZH. Native structure of mosquito salivary protein uncovers domains relevant to pathogen transmission. Nat Commun. 2023;14(1):899. Epub 2023/02/17. doi: 10.1038/s41467-023-36577-y. PubMed PMID: 36797290; PMCID: PMC9935623.

17. Rosenberg J, Ischebeck T, Commichau FM. Vitamin B6 metabolism in microbes and approaches for fermentative production. Biotechnol Adv. 2017;35(1):31–40. Epub 2016/11/29. doi: 10.1016/j.biotechadv.2016.11.004. PubMed PMID: 27890703.

18. Novikova IV, Zhou M, Du C, Parra M, Kim DN, VanAernum ZL, Shaw JB, Hellmann H, Wysocki VH, Evans JE. Tunable Heteroassembly of a Plant Pseudoenzyme-Enzyme Complex. ACS Chem Biol. 2021;16(11):2315–25. Epub 2021/09/15. doi: 10.1021/acschembio.1c00475. PubMed PMID: 34520180; PMCID: PMC9979268.

19. Barra ALC, Ullah N, Morao LG, Wrenger C, Betzel C, Nascimento AS. Structural Dynamics and Perspectives of Vitamin B6 Biosynthesis Enzymes in Plasmodium: Advances and Open Questions. Front Cell Infect Microbiol. 2021;11:688380. Epub 2021/07/31. doi: 10.3389/fcimb.2021.688380. PubMed PMID: 34327152; PMCID: PMC8313854.

20. Rodrigues MJ, Windeisen V, Zhang Y, Guedez G, Weber S, Strohmeier M, Hanes JW, Royant A, Evans G, Sinning I, Ealick SE, Begley TP, Tews I. Lysine relay mechanism coordinates intermediate transfer in vitamin B6 biosynthesis. Nat Chem Biol. 2017;13(3):290–4. Epub 2017/01/17. doi: 10.1038/nchembio.2273. PubMed PMID: 28092359; PMCID: PMC6078385.

21. Matsuura A, Yoon JY, Yoon HJ, Lee HH, Suh SW. Crystal structure of pyridoxal biosynthesis lyase PdxS from Pyrococcus horikoshii. Mol Cells. 2012;34(4):407–12. Epub 2012/10/30. doi: 10.1007/s10059-012-0198-8. PubMed PMID: 23104439; PMCID: PMC3887772.

22. Manzoku M, Ebihara, A., Yokoyama, S., Kuramitsu, S., RIKEN Structural Genomics/Proteomics Initiative (RSGI). Crystal structure of pyridoxine biosynthesis protein from Methanocaldococcus jannaschii 2007. Epub 6 May 2007. doi: 10.2210/pdb2yzr/pdb.

23. Hanford MJ, Peeples TL. Archaeal tetraether lipids: unique structures and applications. Appl Biochem Biotechnol. 2002;97(1):45–62. Epub 2002/03/20. doi: 10.1385/abab:97:1:45. PubMed PMID: 11900115.

24. Arbing MA, Chan S, Shin A, Phan T, Ahn CJ, Rohlin L, Gunsalus RP. Structure of the surface layer of the methanogenic archaean Methanosarcina acetivorans. Proc Natl Acad Sci U S A. 2012;109(29):11812–7. Epub 2012/07/04. doi: 10.1073/pnas.1120595109. PubMed PMID: 22753492; PMCID: PMC3406845.

25. Wagner A, Whitaker RJ, Krause DJ, Heilers JH, van Wolferen M, van der Does C, Albers SV. Mechanisms of gene flow in archaea. Nat Rev Microbiol. 2017;15(8):492–501. Epub 2017/05/16. doi: 10.1038/nrmicro.2017.41. PubMed PMID: 28502981.

26. Deppenmeier U, Johann A, Hartsch T, Merkl R, Schmitz RA, Martinez-Arias R, Henne A, Wiezer A, Baumer S, Jacobi C, Bruggemann H, Lienard T, Christmann A, Bomeke M, Steckel S, Bhattacharyya A, Lykidis A, Overbeek R, Klenk HP, Gunsalus RP, Fritz HJ, Gottschalk G. The genome of Methanosarcina mazei: evidence for lateral gene transfer between bacteria and archaea. J Mol Microbiol Biotechnol. 2002;4(4):453–61. Epub 2002/07/20. PubMed PMID: 12125824.

27. Stach K, Stach W, Augoff K. Vitamin B6 in Health and Disease. Nutrients. 2021;13(9). Epub 2021/09/29. doi: 10.3390/nu13093229. PubMed PMID: 34579110; PMCID: PMC8467949.

28. Abad AND, Seshadri K, Ohashi M, Delgadillo DA, de Moraes LS, Nagasawa KK, Liu M, Johnson S, Nelson HM, Tang Y. Discovery and Characterization of Pyridoxal 5’-Phosphate-Dependent Cycloleucine Synthases. J Am Chem Soc. 2024;146(21):14672–84. Epub 2024/05/15. doi: 10.1021/jacs.4c02142. PubMed PMID: 38743881.

29. di Salvo ML, Safo MK, Contestabile R. Biomedical aspects of pyridoxal 5’-phosphate availability. Front Biosci (Elite Ed). 2012;4(3):897–913. Epub 2011/12/29. doi: 10.2741/E428. PubMed PMID: 22201923.

30. Zhang P, Tsuchiya K, Kinoshita T, Kushiyama H, Suidasari S, Hatakeyama M, Imura H, Kato N, Suda T. Vitamin B6 Prevents IL-1beta Protein Production by Inhibiting NLRP3 Inflammasome Activation. J Biol Chem. 2016;291(47):24517–27. Epub 2016/10/14. doi: 10.1074/jbc.M116.743815. PubMed PMID: 27733681; PMCID: PMC5114405.

31. Wan Z, Zheng J, Zhu Z, Sang L, Zhu J, Luo S, Zhao Y, Wang R, Zhang Y, Hao K, Chen L, Du J, Kan J, He H. Intermediate role of gut microbiota in vitamin B nutrition and its influences on human health. Front Nutr. 2022;9:1031502. Epub 2022/12/31. doi: 10.3389/fnut.2022.1031502. PubMed PMID: 36583209; PMCID: PMC9792504.

32. Sowers KR, Boone JE, Gunsalus RP. Disaggregation of Methanosarcina spp. and Growth as Single Cells at Elevated Osmolarity. Appl Environ Microbiol. 1993;59(11):3832–9. Epub 1993/11/01. doi: 10.1128/aem.59.11.3832-3839.1993. PubMed PMID: 16349092; PMCID: PMC182538.

33. Mastronarde DN. SerialEM: A Program for Automated Tilt Series Acquisition on Tecnai Microscopes Using Prediction of Specimen Position. Microscopy and Microanalysis. 2003;9(S02):1182–3. Epub 24 July 2003. doi: 10.1017/S1431927603445911.

34. Punjani A, Rubinstein JL, Fleet DJ, Brubaker MA. cryoSPARC: algorithms for rapid unsupervised cryo-EM structure determination. Nat Methods. 2017;14(3):290–6. Epub 2017/02/07. doi: 10.1038/nmeth.4169. PubMed PMID: 28165473.

35. Bepler T, Morin A, Rapp M, Brasch J, Shapiro L, Noble AJ, Berger B. Positive-unlabeled convolutional neural networks for particle picking in cryo-electron micrographs. Nat Methods. 2019;16(11):1153–60. Epub 2019/10/09. doi: 10.1038/s41592-019-0575-8. PubMed PMID: 31591578; PMCID: PMC6858545.

36. Croll TI. ISOLDE: a physically realistic environment for model building into low-resolution electron-density maps. Acta Crystallogr D Struct Biol. 2018;74(Pt 6):519–30. Epub 2018/06/07. doi: 10.1107/S2059798318002425. PubMed PMID: 29872003; PMCID: PMC6096486.

37. Meng EC, Goddard TD, Pettersen EF, Couch GS, Pearson ZJ, Morris JH, Ferrin TE. UCSF ChimeraX: Tools for structure building and analysis. Protein Sci. 2023;32(11):e4792. Epub 2023/09/29. doi: 10.1002/pro.4792. PubMed PMID: 37774136; PMCID: PMC10588335.

38. Liebschner D, Afonine PV, Baker ML, Bunkoczi G, Chen VB, Croll TI, Hintze B, Hung LW, Jain S, McCoy AJ, Moriarty NW, Oeffner RD, Poon BK, Prisant MG, Read RJ, Richardson JS, Richardson DC, Sammito MD, Sobolev OV, Stockwell DH, Terwilliger TC, Urzhumtsev AG, Videau LL, Williams CJ, Adams PD. Macromolecular structure determination using X-rays, neutrons and electrons: recent developments in Phenix. Acta Crystallogr D Struct Biol. 2019;75(Pt 10):861–77. Epub 2019/10/08. doi: 10.1107/S2059798319011471. PubMed PMID: 31588918; PMCID: PMC6778852.

39. Berman H, Henrick K, Nakamura H. Announcing the worldwide Protein Data Bank. Nat Struct Biol. 2003;10(12):980. Epub 2003/11/25. doi: 10.1038/nsb1203-980. PubMed PMID: 14634627.

40. Madeira F, Madhusoodanan N, Lee J, Eusebi A, Niewielska A, Tivey ARN, Lopez R, Butcher S. The EMBL-EBI Job Dispatcher sequence analysis tools framework in 2024. Nucleic Acids Res. 2024. Epub 2024/04/10. doi: 10.1093/nar/gkae241. PubMed PMID: 38597606.

41. Robert X, Gouet P. Deciphering key features in protein structures with the new ENDscript server. Nucleic Acids Res. 2014;42(Web Server issue):W320-4. Epub 2014/04/23. doi: 10.1093/nar/gku316. PubMed PMID: 24753421; PMCID: PMC4086106.

42. UniProt C. UniProt: the Universal Protein Knowledgebase in 2023. Nucleic Acids Res. 2023;51(D1):D523–D31. Epub 2022/11/22. doi: 10.1093/nar/gkac1052. PubMed PMID: 36408920; PMCID: PMC9825514.

43. Mooney S, Leuendorf JE, Hendrickson C, Hellmann H. Vitamin B6: a long known compound of surprising complexity. Molecules. 2009;14(1):329–51. Epub 2009/01/16. doi: 10.3390/molecules14010329. PubMed PMID: 19145213; PMCID: PMC6253932.

44. Oka T. Modulation of gene expression by vitamin B6. Nutr Res Rev. 2001;14(2):257–66. Epub 2001/12/01. doi: 10.1079/NRR200125. PubMed PMID: 19087426.

